# Decreased alertness reconfigures cognitive control networks

**DOI:** 10.1101/831727

**Authors:** Andres Canales-Johnson, Lola Beerendonk, Salome Blain, Shin Kitaoka, Alejandro Ezquerro-Nassar, Stijn Nuiten, Johannes Fahrenfort, Simon van Gaal, Tristan A. Bekinschtein

## Abstract

Humans’ remarkable capacity to flexibly adapt their behaviour based on rapid situational changes is termed cognitive control. Intuitively, cognitive control is thought to be affected by the state of alertness, for example, when drowsy we feel less capable of adequately implementing effortful cognitive tasks. Although scientific investigations have focused on the effects of sleep deprivation and circadian time, little is known about how natural daily fluctuations in alertness in the regular awake state affect cognitive control. Here we combined a conflict task in the auditory domain with EEG neurodynamics to test how neural and behavioural markers of conflict processing are affected by fluctuations in alertness. Using a novel computational method, we segregated alert and drowsy trials from two testing sessions and observed that, although participants (both sexes) were generally sluggish, the typical Conflict Effect reflected in slower responses to conflicting information compared to non-conflicting information was still intact, as well as the moderating effect of previous conflict (Conflict Adaptation). However, the typical neural markers of cognitive control-local midfrontal-theta band power changes-that participants show during full alertness were no longer noticeable when alertness decreased. Instead, when drowsy, we found an increase in long-range information sharing (connectivity) between brain regions in the same frequency band. These results show the resilience of the human cognitive control system when affected by internal fluctuations of alertness, and suggest neural compensatory mechanisms at play in response to physiological pressure during diminished alertness.

**Significance Statement:** The normal variability in alertness we experience in daily tasks is rarely taking into account in cognitive neuroscience. Here we studied neurobehavioral dynamics of cognitive control with decreasing alertness. We used the classic Simon Task where participants hear the word “left” or “right” in the right or left ear, eliciting slower responses when the word and the side are incongruent - the conflict effect. Participants performed the task both while fully awake and while getting drowsy, allowing for the characterisation of alertness modulating cognitive control. The changes in the neural signatures of conflict from local theta oscillations to a long-distance distributed theta network suggests a reconfiguration of the underlying neural processes subserving cognitive control when affected by alertness fluctuations.

## INTRODUCTION

Cognitive control is the capacity of making quick adjustments to cognitive processes in order to optimally solve the task at hand. One proposed mechanism involves allocating attention to task-relevant information and ignoring non-relevant, sometimes conflictive, information (Desimone and Duncan, 1995; Miller and Cohen, 2001; Egner and Hirsch, 2005). The ability to deal with conflicting information is often studied using “conflict tasks”, which typically induce response (or stimulus) conflict by triggering an automatic response that has to be overcome to decide correctly (e.g. Stroop/Simon tasks). For example, when a Dutch person drives in England, they must override the automatic tendency to turn right on a roundabout, and go left instead. Experiencing these types of conflict has shown to increase the level of cognitive control on the next occasion, when encountering a similar conflicting situation. This process-termed conflict adaptation-seems necessary to smooth future decisions and avoid further mistakes (Gratton et al., 1992). Here we combine a behavioural conflict task with electroencephalography (EEG) to study the modulatory effect of arousal fluctuations on decision-making in the face of conflict.

Alertness, as an arousal component, is controlled at various levels in the mammalian brain, including the pons and midbrain, upper brainstem and thalamus as well as cortical interactions (Bekinschtein et al., 2009). Arousal is dependent on circadian factors and sleep pressure (Borbély et al., 2016) and in turn affects the efficiency of cognitive processes. How levels of wakefulness modulate attentional processes and cognitive control is commonly studied in sleep deprivation and circadian cycle studies, but less often during normal waking fluctuations (Goupil and Bekinschtein, 2012). Both sleep deprivation and drops in circadian time lead to cognitive performance decrements (Tucker et al., 2010; Ratcliff and Van Dongen, 2011; Gunzelmann et al., 2012) but surprisingly, the performance modulation imposed by changes in wakefulness on complex tasks appears to be less severe than their effects on simple tasks (Harrison et al., 2000). Specifically, studies focusing on (cognitive/response) conflict have failed to indicate increased interference effects with sleep deprivation and circadian time (Sagaspe et al., 2006; Cain et al., 2011; Bratzke et al., 2012), but consistently show overall slower responses during increased sleepiness or lower arousal. However, Gevers et al. (2015) recently uncovered an interesting dissociation, although conflict effects on the current trial did not seem to change after a night of sleep deprivation, across trial conflict adaptation effects did. These results nicely converge with studies on the relationship between conflict awareness and conflict processing, as conflict detection seems much less dependent on conflict experience than conflict adaptation (van Gaal et al., 2010; Jiang et al., 2015), suggesting that conflict detection is more automatic-less effortful-than conflict adaptation.

Fluctuations in cognitive control are shown to be associated with changes in activity patterns in the medial frontal cortex (MFC) and the dorsolateral prefrontal cortex (DLPFC) (Robbins, 1996; Swick et al., 2011; Gläscher et al., 2012; Cai et al., 2016). In EEG recordings, conflict-related processes are often measured by quantifying the power of theta-band neural oscillations (4-8 Hz) (Luu et al., 2004; Trujillo and Allen, 2007; Cohen et al., 2008; Cavanagh et al., 2010; Nigbur et al., 2012; Cohen and van Gaal, 2014). In combination with a recently validated method to automatically detect drowsiness periods from EEG (Jagannathan et al., 2018) we here use conflictive information to map behavioural and neural markers of cognitive control as they get modulated by ongoing fluctuations in arousal.

## METHODS

### Participants

Thirty-three healthy human participants (18 female) aged 18 to 30 (M=23.1, SD=2.8), recruited from the University of Cambridge (Cambridge, United Kingdom), participated in this experiment for monetary compensation. All participants had normal or corrected-to-normal vision and had no history of head injury or physical and mental illness. This study was approved by the local ethics committee of the University of Cambridge and written informed consent was obtained from all participants after explanation of the experimental protocol.

### Experimental task

Participants performed an auditory version of the Stroop task (Stroop, 1935). Recorded samples of a native speaker saying “left” or “right” were presented to participants’ left or right ear through ear buds, resulting in four types of stimuli (i.e. “left” in left ear, “left” in right ear, “right” in right ear, “right” in left ear). Stimuli were congruent when the word meaning corresponded to its physical location (e.g. left in left ear) and incongruent otherwise (e.g. “left” in right ear). All four types of stimuli were presented equally often, but in a random order. Participants were asked to report the location depicted by the voice (i.e. word meaning; the words left or right), while ignoring its physical location (i.e. left or right ear) by pressing one of two buttons on a response box. There was no practice block and no feedback on performance throughout the task. The time between a response and the following stimulus varied randomly between 2 and 2.5 seconds. The inter stimulus interval was fixed to 2 seconds in the absence of a response within that time frame. As a result, the inter stimulus interval could vary from 2 seconds (response absent) to 4.49 seconds (maximum response latency of 1.99 seconds + maximum response stimulus interval of 2.5).

### Procedure

Participants were instructed to get a normal night’s rest on the night previous to testing. Testing started between 9 am and 5 pm and lasted approximately 3 hours. Upon arrival at the testing room, participants were sat down in a comfortable adjustable chair in an electrically shielded room. Participants were fitted with an EGI electrolyte 129-channel cap (Electrical Geodesics, Inc. systems) after receiving the task instructions and subsequently signing the informed consent. Task instructions were to respond as fast and accurate as possible, to keep bodily movements to a minimum and to keep the eyes closed throughout the experiment. Participants were asked to report their answers with their thumbs (i.e. left thumb for the word ‘left’ and vice versa) on two buttons of a four-button response box that rested on their lap or abdomen. In the first part of the session, participants were instructed to stay awake with their eyes closed whilst performing the task. The back of the chair was set up straight and the lights in the room were on. This part of the experiment lasted for 500 trials and lasted for approximately 25 minutes. Right afterwards, the task was performed while participants were allowed to fall asleep. The chair was reclined to a comfortable position, the lights were turned off and participants were offered a pillow and blanket. Participants were told that the experimenter would wake them up by making a sound (i.e. knocking on desk or wall) if they missed 5 consecutive trials. This procedure prevents participants taking extended naps during the session as it takes only 1 responsive trial (following an unresponsive one) to restart the count. In this way, a session can accumulate several unresponsive trials in the absence of extended unresponsive periods. This part of the experiment lasted for 2000 trials and lasted for approximately 1.5 hours. At the end of the session, participants were sat upright and the EEG cap was removed. Stimuli were presented using PsychToolbox software on a Mac computer and data were acquired using NetStation software (Electrical Geodesics, Inc. Systems) on another Mac computer.

For each individual session (awake and drowsy), we performed an automatic alertness classification in order to sub-select the awake trials of the awake session, and the drowsy trials of the drowsy one (see below). As a result of the automatic classification, 93% of the trials from the awake session were classified as awake, and selected for further analyses while discarding the rest (average across 33 participants). Similarly, for the drowsy condition, 62% of the trials were classified as drowsy, selected for further analyses, and discarding the rest.

### Wakefulness classification

The automatic classification of alertness levels involved classifying periods of the experimental session into ‘awake’ and ‘drowsy’. The pre-trial period (−1500 to 0ms) before each tone was used in classifying the corresponding trial as awake or drowsy. Pre-trial epochs were analysed using the micro-measures algorithm (Jagannathan et al., 2018) and each trial was classified as ‘alert’, ‘drowsy (mild)’ or ‘drowsy (severe)’.

The micro-measures algorithm computes predictor variance and coherence features on the pre-trial period, and subsequently classifies trials as alert and drowsy using Support Vector Machine (SVM). Predictor variance is computed in the occipital electrodes (O1, Oz, O2) in different frequency bands (band 1: 2-4 Hz; band 2: 8-10 Hz, band 3: 10-12 Hz; band 4: 2-6 Hz), and coherence is computed across selected electrodes in the occipital (O1, Oz, O2), frontal (F7, F8, Fz), central (C3, C4;), and temporal (T7, T8, TP8, FT10, TP10) regions in the following bands (delta: 1-4 Hz; alpha: 7-12 Hz; sigma: 12-16 Hz; and gamma: 16-30 Hz). Intuitively, while predictor variance captures the variance in the signal explained by different frequency bands, coherence captures the spectral correlation across electrodes by frequency band. SVM later on uses these features for alertness classification, where pre-trails containing 100% and >50% of alpha oscillations are classified as ‘awake’, and trials containing 50% of alpha oscillations, EEG flattening, and ripples are classified as ‘drowsy mild’. Finally, further detectors are used to further sub-classify them into ‘drowsy severe’ (vertex waves, k-complex and spindles).

To select true alert trials from the ‘awake’ condition, we used only trials from the alert blocks and removed all those marked as ‘drowsy’ (purple in Figure 1C). Similarly, ‘drowsy (mild)’ and ‘drowsy (severe)’ from the drowsy blocks were selected as true drowsy trials during the drowsy condition (green in Figure 1C). The periods where the trials were classified as ‘drowsy (mild)’ and ‘drowsy (severe) corresponded to a higher degree of misses (time-out responses) further confirming, behaviourally, the low alertness of these sections of the drowsy session. The total number of trials across the 33 participants was 26045 for the ‘awake’ and 33306 for the ‘drowsy’ conditions.

**Figure 1.**
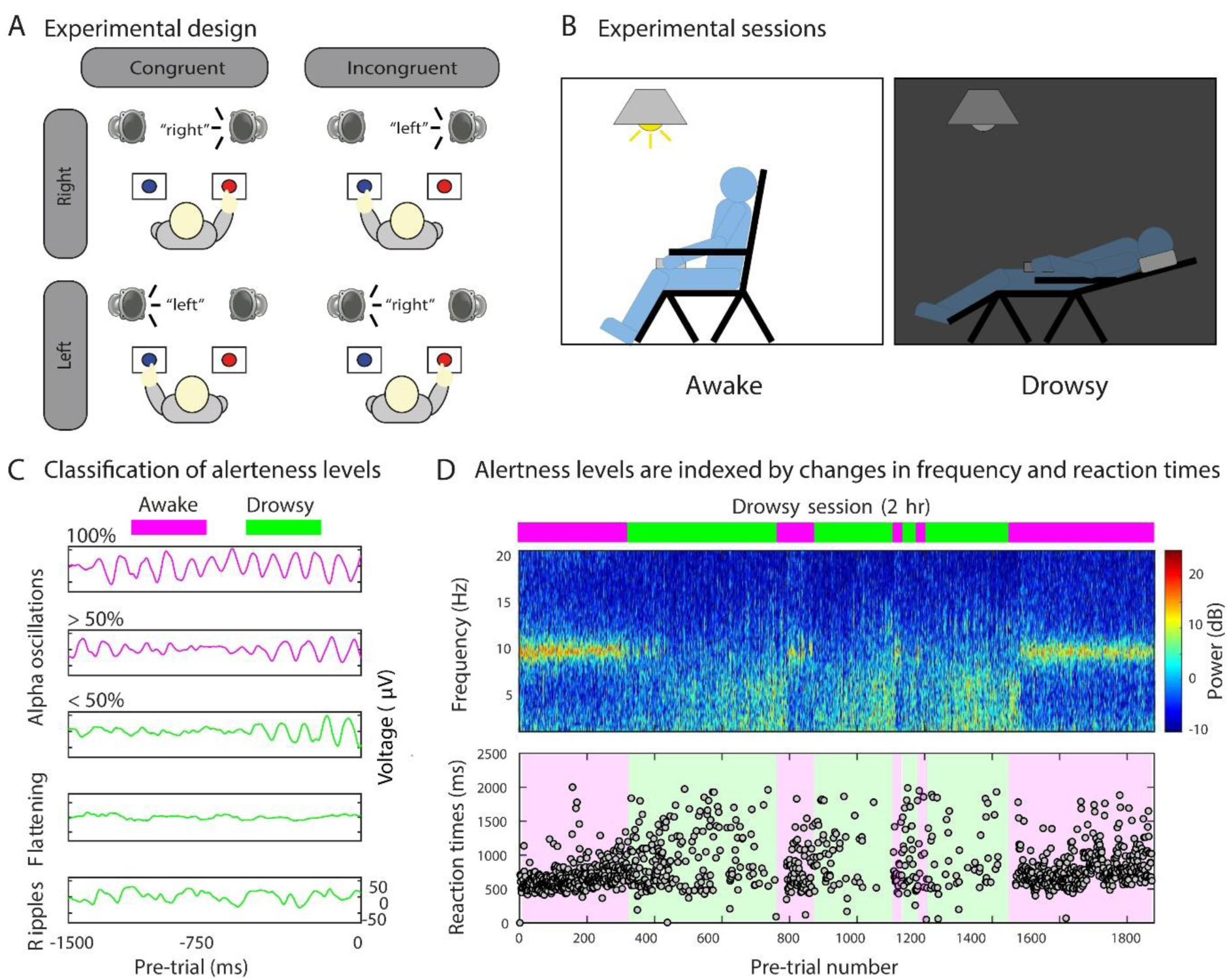
Experimental paradigm and alertness level classification. **(A)** Schematic representation of the experimental design. Participants were instructed to report the semantics (“left” or “right”) of an auditory stimulus via a button press with their left or right hand, respectively, and to ignore the spatial location at which the auditory stimulus was presented. Sound content of the auditory stimuli could be congruent or incongruent with its location of presentation (50% congruent/incongruent trials). **(B)** Schematic representation of the experimental sessions. In the awake session participants were instructed to stay awake with their eyes closed whilst performing the task with the back of the chair set up straight and the lights on. Immediately after, in the drowsy session, the task was performed while participants were allowed to fall asleep with their chair reclined to a comfortable position and the lights off. **(C)** Automatic classification of alertness levels. For each session (awake and drowsy), pre-trial periods (−1500 to 0 ms) were used for defining awake (purple) and drowsy (green) trials. Pre-trials containing 100% and >50% of alpha oscillations were classified as ‘awake’. Similarly, pre-trials containing <50% of alpha oscillations, EEG flattening, ripples, and other grapho-elements were classified as ‘drowsy’ (see Methods for details). Thus, only the trials classified as ‘awake’ from the awake session, and those classified as ‘drowsy’ from the drowsy session were sub-selected for further analyses **(D)** *Upper panel.* Automatic classification of alertness during a drowsy session (representative participant, occipital electrode). The frequency profile depicts changes in the power level in different bands during the pre-trial period, and the bars on top represent pre-trials classified as awake (purple) or drowsy (green). *Lower panel.* The variability in the reaction times (lower panel) closely follows the changes in the frequency profile (upper panel) from alpha (higher RT variability in green) to theta (lower RT variability in purple) obtained using the pre-trial information.

### Rationale for conflict effect and conflict adaptation effect analyses

Although the proportion of missed trials due to unresponsiveness was higher for the drowsy condition (42.80%) than for the awake condition (13.85%) (t_32_=-9.75; p<0.001), the available number of trials for further analyses was comparable between conditions (overall number of trials accumulated across subjects; awake: 22398 trials [median between participants: 698; range: 524-954]; drowsy: 19051 trials [median between participants: 923; range: 790-1278]). However, the situation changed for the conflict adaptation analysis. The fact that conflict adaptation is evaluated over sequences of two consecutive responded trials (e.g. an incongruent trial preceded by a congruent one) brought as a consequence a greater reduction in the number of trials in the drowsy state (12008 out of 19051 trials, 46.38% reduction) as compared to awake trials (20138 out of 22398 trials, 10.09% reduction). Therefore, for the conflict effect analyses in RT and theta-power, the (RM) ANOVA incorporated only two factors in order to maximize the number of trials (and therefore statistical power) within conditions and to balance the proportion of trials between conditions (alertness: awake vs drowsy; and congruency: congruent vs incongruent). In contrast, for the conflict adaptation analyses (RT and theta-power), three factors were considered for the (RM) ANOVA: alertness (awake versus drowsy), previous-trial congruency (congruent vs incongruent) and current-trial congruency (congruent vs incongruent).

### Behavioral data analysis

The first trial of every block, incorrect or missed trials, trials following incorrect responses and trials with an RT<200 ms were excluded from behavioral analyses. Conflict on trial *n* has been found to cause increased error rates (ERs) and prolonged reaction times (RTs), as compared to when no conflict is present. This current trial effect of conflict can be modulated by previously experienced conflict on trial n-1, a phenomenon called conflict adaptation. In order to investigate whether current trial conflict effects and the modulation thereof by previous conflict were present, we performed repeated measures (RM) ANOVA on ERs and RTs between alertness (awake, drowsy), current trial congruency (congruent, incongruent) and previous trial congruency (congruent, incongruent). Additional post-hoc (RM) ANOVA, awake and drowsy conditions separately were performed. In case of null-findings, we applied a Bayesian repeated measures ANOVA with similar factors, to verify if there is actual support of the null-hypothesis. We also performed such Bayesian ANOVAs for any null-findings in our EEG data.

### EEG recordings and pre-processing

EEG signals were recorded with 128-channel HydroCel Sensors using a GES300 Electrical Geodesic amplifier at a sampling rate of 500 Hz using the NetStation software. During recording and analyses, the electrodes’ average was used as the reference electrode. Two bipolar derivations were designed to monitor vertical and horizontal ocular movements. Following Chennu et al (2014), data from 93 channels over the scalp surface were retained for further analysis. Channels on the neck, cheeks and forehead, which reflected more movement-related noise than signal, were excluded. Continuous EEG data was epoched from −1500 to 2000 ms around stimulus onset. Eye movement contamination (blinks were rare as eyes were closed, vertical and horizontal saccades or slow movements were also infrequent), muscle artefacts (i.e. cardiac and neck movements) were removed from data before further processing using an independent component analysis (ICA) (Delorme and Makeig 2004). All conditions yielded at least 96% of artefact-free trials. Trials that contained voltage fluctuations exceeding ± 200 μV, transients exceeding ± 100 μV were removed. No low-pass or high-pass filtering was performed during the pre-processing stage. The EEGLAB MATLAB toolbox was For each individual session adopted as it can significantly affect classification scores during MVPA analysis (Driel et al., 2019). Thus, we applied a function designed for rejecting abnormal slow-frequency trends mainly caused by artifactual eye-related currents. To detect such drifts, we used a detrending function implemented in EEGLAB that fits the data to a straight line and marks the trial for rejection if the slope exceeds a given threshold. The slope is expressed in microvolt over the whole epoch (50, for instance, would correspond to an epoch in which the straight-line fit value might be 0 μv at the beginning of the trial and 50 μ v at the end). The minimal fit between the EEG data and a line of minimal slope is determined using a standard R-square measure.

### EEG time-frequency analysis

Epochs were grouped based on current and previous trial congruency, creating four trial conditions. Then, EEG-traces were decomposed into time-frequency charts from 2 Hz to 30 Hz in 15 linearly spaced steps (2 Hz per bin). The power spectrum of the EEG-signal (as obtained by the fast Fourier transform) was multiplied by the power spectra of complex Morlet wavelets 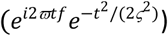 with logarithmically spaced cycle sizes ranging from 3 to 12. The inverse Fourier transform was then used to acquire the complex signal, which was converted to frequency-band specific power by squaring the result of the convolution of the complex and real parts of the signal (*real*[*Z(t)*]^2^ + *imag[Z(t)]*^2^). The resulting time-frequency data were then averaged per subject and trial type. Finally, time-frequency traces were transformed to decibels (dB) and normalized to a baseline of −400ms to −100 ms before stimulus onset, according to: 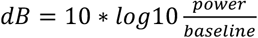(Cohen and van Gaal, 2014).

We tested the hypothesis that midfrontal theta-power would increase following the presentation of conflicting stimuli according to previous literature (Nigbur et al., 2012; Cohen and Ridderinkhof, 2013; Pastötter et al., 2013; Cohen and van Gaal, 2014). Therefore, we selected electrodes in a fronto-central spatial region of interest (ROI) to run our analyses (Figure 3). In order to find a time-frequency ROI for subsequent analyses in the spectral and information-theory domain, data from within the spatial ROI were averaged across the awake and drowsy experimental sessions for congruent and incongruent trials, separately. Next, current trial conflict was calculated (I-C) for all participants.

To test for significant time-frequency ROI in which overall conflict was present (Figure 3A), a cluster-based nonparametric statistical test implemented in FieldTrip (Maris and Oostenveld, 2007) was used. In brief, time-frequency charts (−200 to 1200 ms) were compared in pairs of experimental conditions (incongruent vs. congruent). For each such pairwise comparison, epochs in each condition were averaged subject-wise. These averages were passed to the analysis procedure of FieldTrip, the details of which are described elsewhere (Maris and Oostenveld, 2007). In short, this procedure compared corresponding temporal points in the subject-wise averages using independent samples t-tests for between-subject comparisons. Although this step was parametric, FieldTrip uses a nonparametric clustering method to address the multiple comparisons problem. t values of adjacent temporal points whose P values were lower than 0.05 were clustered together by summating their t values, and the largest such cluster was retained. This whole procedure, i.e., calculation of t values at each temporal point followed by clustering of adjacent t values, was then repeated 1000 times, with recombination and randomized resampling of the subject-wise averages before each repetition. This Monte Carlo method generated a nonparametric estimate of the p-value representing the statistical significance of the originally identified cluster. The cluster-level t value was calculated as the sum of the individual t values at the points within the cluster.

Then, time-frequency power was extracted from this ROI for each participant and used as input for (RM) ANOVAs between alertness (awake, drowsy) and congruency (congruent, incongruent) for the conflict effect analysis. Subsequently, separate (RM) ANOVA were performed on the same ROI data for post-hoc inspection of significant effects for conflict adaptation between alertness (awake, drowsy), previous-trial congruency (congruent, incongruent) and current-trial congruency (congruent, incongruent). Finally, separate (RM) ANOVAs for the awake and drowsy conditions were performed on the same ROI data for post-hoc inspection of significant effects for conflict adaptation (current trial congruency vs previous trial congruency).

### EEG source reconstruction

To visualize the brain origins of the univariate conflict effect, cortical sources of subject-wise averaged time-frequency charts within the theta-band ROI (Figure 3) were reconstructed using Brainstorm (Tadel et al., 2011). The forward model was calculated using the OpenMEEG Boundary Element Method (Gramfort et al., 2010) on the cortical surface of a template MNI brain (colin27) with 1 mm resolution. The inverse model was constrained using weighted minimum-norm estimation (Baillet et al., 2001) to calculate source activation. To plot cortical maps, grand-averaged activation values were baseline corrected by z-scoring the baseline period (−400 to −100 ms window) to each time point, and spatially smoothed with a 5-mm kernel. This procedure was applied separately for the overall, awake and drowsy conflict effect.

### EEG multivariate spectral decoding

In addition to the univariate approach, a multivariate spectral decoding model was applied on the time-frequency data. This was done both because of the higher sensitivity of multivariate analyses, and well as to inspect if and to what extent different stimulus features (i.e. location and sound content) were processed in awake and drowsy conditions. The ADAM-toolbox was used on raw EEG data, that was transformed to time-frequency using default methods but with similar settings epochs: −200 ms to 1200 ms, 2Hz-30Hz (Fahrenfort et al., 2018). As unbalanced designs (i.e. asymmetrical trial counts between conditions) can have a number of unintended or biased effects on the conclusion that can be drawn from the analysis, we performed a balanced decoding between conditions per state of alertness (Figure 3). Balancing the number of trials between conditions has been shown to convey clear performance benefits for linear discriminant analysis and area under the curve, which are the classification algorithm and default performance metric that ADAM uses.

Trials were classified according to current trial stimulus content (i.e. sound location and sound content) resulting in 4 trial types. Note that this is different from the univariate analyses, where trials were classified according to current and previous trial conflict. As decoding algorithms are known to be time-consuming, data were resampled to 64Hz. Next, a backward decoding algorithm, using either stimulus location, stimulus sound contents or congruency as stimulus class, was applied according to a ten-fold cross-validation scheme. A linear discriminant analysis (LDA) was used to discriminate between stimulus classes (e.g. left versus right ear bud location etc.) after which classification accuracy was computed as the area under the curve (AUC), a measure derived from Signal Detection Theory. AUC scores were tested per time-point with double-sided t-tests across participants against a 50% chance-level. These t-tests were corrected for multiple comparisons over time, using cluster-based permutation tests (p<0.05, 1000 iterations). This procedure yields time clusters of significant above-chance classifier accuracy, indicative of information processing. Note that this procedure yields results that should be interpreted as fixed effects (Allefeld et al., 2016), but is nonetheless standard in the scientific community.

### Information sharing analysis: weighted symbolic mutual information (wSMI)

In order to quantify the information sharing between electrodes we computed the weighted symbolic mutual information (wSMI) (King et al., 2013; Sitt et al., 2014; Imperatori et al., 2019). It assesses the extent to which the two signals present joint non-random fluctuations, suggesting that they share information. wSMI has three main advantages: (i) it allows for a rapid and robust estimation of the signals’ entropies; (ii) it provides an efficient way to detect non-linear coupling; and (iii) it discards the spurious correlations between signals arising from common sources, favouring non-trivial pairs of symbols. For each trial, wSMI is calculated between each pair of electrodes after the transformation of the EEG signals into sequence of discrete symbols defined by the ordering of k time samples separated by a temporal separation τ. The symbolic transformation depends on a fixed symbol size (k = 3, that is, 3 samples represent a symbol) and a variable τ between samples (temporal distance between samples) which determines the frequency range in which wSMI is estimated. In our case, we chose τ = 32 to specifically isolate wSMI in theta-band. The frequency specificity *f* of wSMI is related to k and τ as:

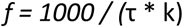

As per the above formula, with a kernel size k of 3, τ values of 32 ms hence produced a sensitivity to frequencies below 10 Hz with and spanning the theta-band (~4-9 Hz). Control results were obtained with τ value of 24 ms (alpha-band: ~10-14 Hz) and τ value of 16 ms (beta-band: ~14-20 Hz).

wSMI was estimated for each pair of transformed EEG signals by calculating the joint probability of each pair of symbols. The joint probability matrix was multiplied by binary weights to reduce spurious correlations between signals. The weights were set to zero for pairs of identical symbols, which could be elicited by a unique common source, and for opposite symbols, which could reflect the two sides of a single electric dipole. wSMI is calculated using the following formula:

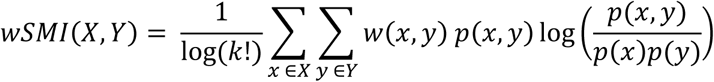

where *x* and *y* are all symbols present in signals *X* and *Y* respectively, w*(x,y)* is the weight matrix and *p(x,y)* is the joint probability of co-occurrence of symbol *x* in signal *X* and symbol *y* in signal *Y*. Finally, *p(x)* and *p(y)* are the probabilities of those symbols in each signal and *K*! is the number of symbols - used to normalize the mutual information (MI) by the signal’s maximal entropy. The time window in which theta-band and alpha-band (control) wSMI was calculated was determined based on the significant time window observed in the spectral contrast of Figure 3A (380-660 ms). Similarly, beta-band wSMI (control) was determined based on the spectral time window of Figure 3A (580ms - 728ms).

#### Statistics

Statistical analyses were performed using MATLAB (2016a), Jamovi (Version 0.8.1.6) [Computer Software] (Retrieved from https://www.jamovi.org) (open source), and JASP Team (2018; JASP; version 0.8.4 software) statistical software. When reported, BF_01_ refers to the Bayes Factor in favour of the null hypothesis.

## RESULTS

While fully awake as well as while becoming drowsy, participants performed an auditory Simon task where they heard the words “left” or “right”, from either the left or right side in space. Participants were instructed to respond according to the meaning of the sound (e.g. “left” requires left-hand response, Figure 1A). We hypothesised an increase in reaction times to all stimuli-a typical marker of drowsiness-but expected that conflict detection mechanisms would remain relatively preserved (in behaviour and theta oscillations), similar to studies showing preserved processing of conflicting information at reduced levels of stimulus awareness (van Gaal et al., 2010; Jiang et al., 2015, 2018). We expected the sharpest decline in performance and conflict processing when focusing on across trial conflict adaptation mechanisms (Jiang et al., 2015), but less so for the current trial conflict effect.

### Behavioural results

#### Conflict effect in reaction times

First, we analysed the reaction times (RT) with the factors alertness conditions (awake, drowsy) and trial congruency (congruent, incongruent). A repeated measures (RM) ANOVA revealed a main effect of alertness (F_1,32_=92.96; p<0.001; 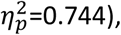, reflecting a slowing of RT during drowsy state compared to awake state. Further, a main effect of congruency (F_1,32_=51.93; p<0.001; 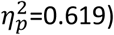 was observed, manifested as slower RTs for incongruent trials compared to congruent trials (“the conflict effect”). The conflict effect was positive for the majority of the participants (30 out of 33 participants). Interestingly, no reliable interaction between alertness and congruency was observed (F_1,32_=0.542; p=0.468; 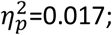 BF_01_=4.199). Next, we focused on the effects for the awake and drowsy conditions separately. Within the awake condition, RTs were slower for incongruent trials compared to congruent trials (main effect of congruency: F_1,32_=59.16; p<0.001; 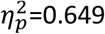 and the effects were positive for the majority of the participants (30 out of 33 participants). A similar conflict effect was observed when participants were drowsy (F_1,32_=9.642; p=0.004; 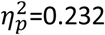 with a positive effect for the majority of the participants (26 out of 33 participants).

#### Conflict adaptation effect in reaction times

In the next analysis, we focused on the conflict adaptation effect, as indicated by a smaller conflict effect when the current trial was preceded by an incongruent trial as compared when it was preceded by a congruent one. A (RM) ANOVA performed on conflict adaptation across alertness levels revealed a conflict adaptation effect (interaction previous x current trial congruency: F_1,32_=29.885; p<0.001; 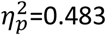, Figure 2C). The conflict adaptation effect was positive for the majority of the participants in the awake (26 out of 33 participants; F_1,32_=9.642; p=0.004; 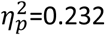, Figure 2A middle) and drowsy (26 out of 33 participants, F_1,32_=7.318; p=0.011; 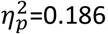, Figure 2D) conditions. Furthermore, no reliable interaction between conflict adaptation and alertness was observed (F_1,32_=0.683; p=0.415; 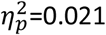 BF_01_=4.199).

**Figure 2.**
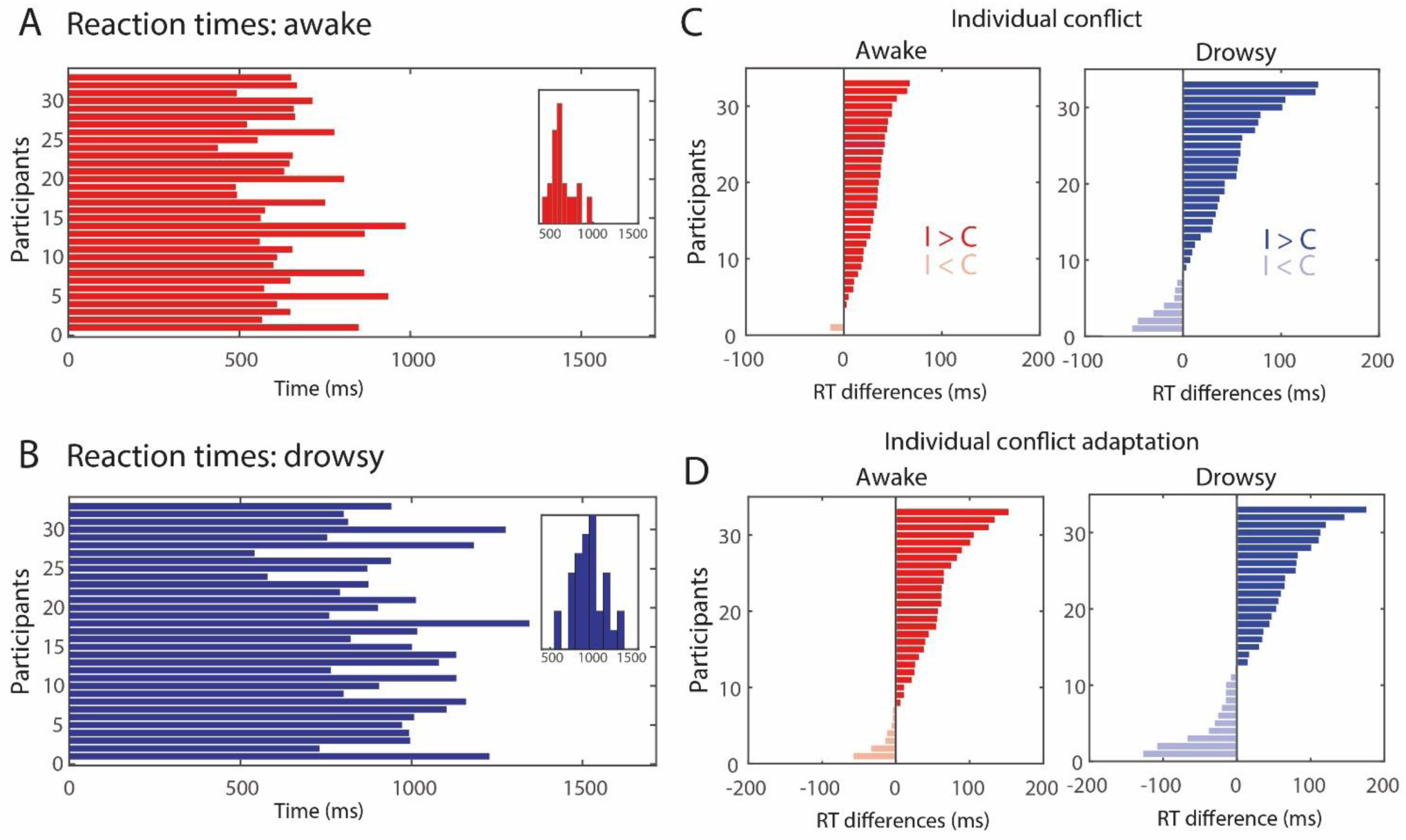
Behavioral results in awake and drowsy. Overall reaction times for the awake **(A)** and drowsy **(B)** conditions. Individual conflict **(C)** and conflict adaptation effects **(D)** for the awake and drowsy conditions in reaction times.

#### Error rates

This version of the Simon Tasks is well tuned to test for conflict effect in reaction times and less for error rates, however, a (RM) ANOVA performed on error rates and alertness conditions (alertness x conflict x conflict adaptation) revealed that participants made more errors during drowsy than during awake (main effect of alertness: F_1,32_=18.29; p<0.001; 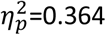. However, conflict (F_1,32_=2.357; p=0.135; 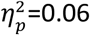 BF_01_=1.24) and conflict adaptation (F_1,32_=0.862; p=0.360; 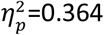 BF_01_=4.14) effects on error rate were not reliable, nor the interaction between conflict adaptation and alertness (F_1,32_=3.177; p=0.084; 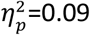 BF_01_=2.35). Finally, when the analyses were performed separately by alertness conditions, the awake state showed a conflict effect (F_1,32_=24.152; p<0.001; 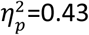 and conflict adaptation F_1,32_=8.567; p=0.006; 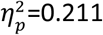 but the drowsy condition did not (conflict: F_1,32_=1.41; p=0.243; 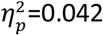 BF_01_=5.149; conflict adaptation: F_1,32_=1.88; p=0.180; 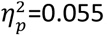 BF_01_=13.685).

Overall, we observed that although participants were generally slower and made more errors during the drowsy condition, the conflict effect reflected in slower responses to conflicting information compared to non-conflicting information was still present, as well as the moderating effect of previous trial conflict observed in the conflict adaptation effect.

### Midfrontal theta-band oscillations and source reconstruction

Upon establishing that conflict and conflict adaptation effects are present in both awake and drowsy states, we proceed to test whether medial frontal (MF) conflict detection processes, typically reflected in short-lived oscillatory dynamics in the theta-band (Nigbur et al., 2012; Cohen and Donner, 2013; Cohen and van Gaal, 2014; Jiang et al., 2015), were present during awake and drowsy states as well. In order to determine the time-frequency cluster for assessing conflict and conflict adaptation effects, we first analysed the overall conflict effect, irrespective of alertness condition or previous trial congruency (I-C, averaged over awake and drowsy sessions) (Figure 3A). Replicating previous studies (Nigbur et al., 2012; Jiang et al., 2015), current trial conflict induced increased theta-band power at MF electrodes (cluster p=0.028; frequency range: 4Hz–8Hz, time range: 250ms–625ms, see encircled region in black, solid line, in Figure 3A). The area within this time-frequency (T-F) cluster was used for follow-up analyses. Next, we tested whether these conflict-related theta-band dynamics in this cluster were modulated by alertness and previous trial congruency, which was indeed the case.

**Figure 3.**
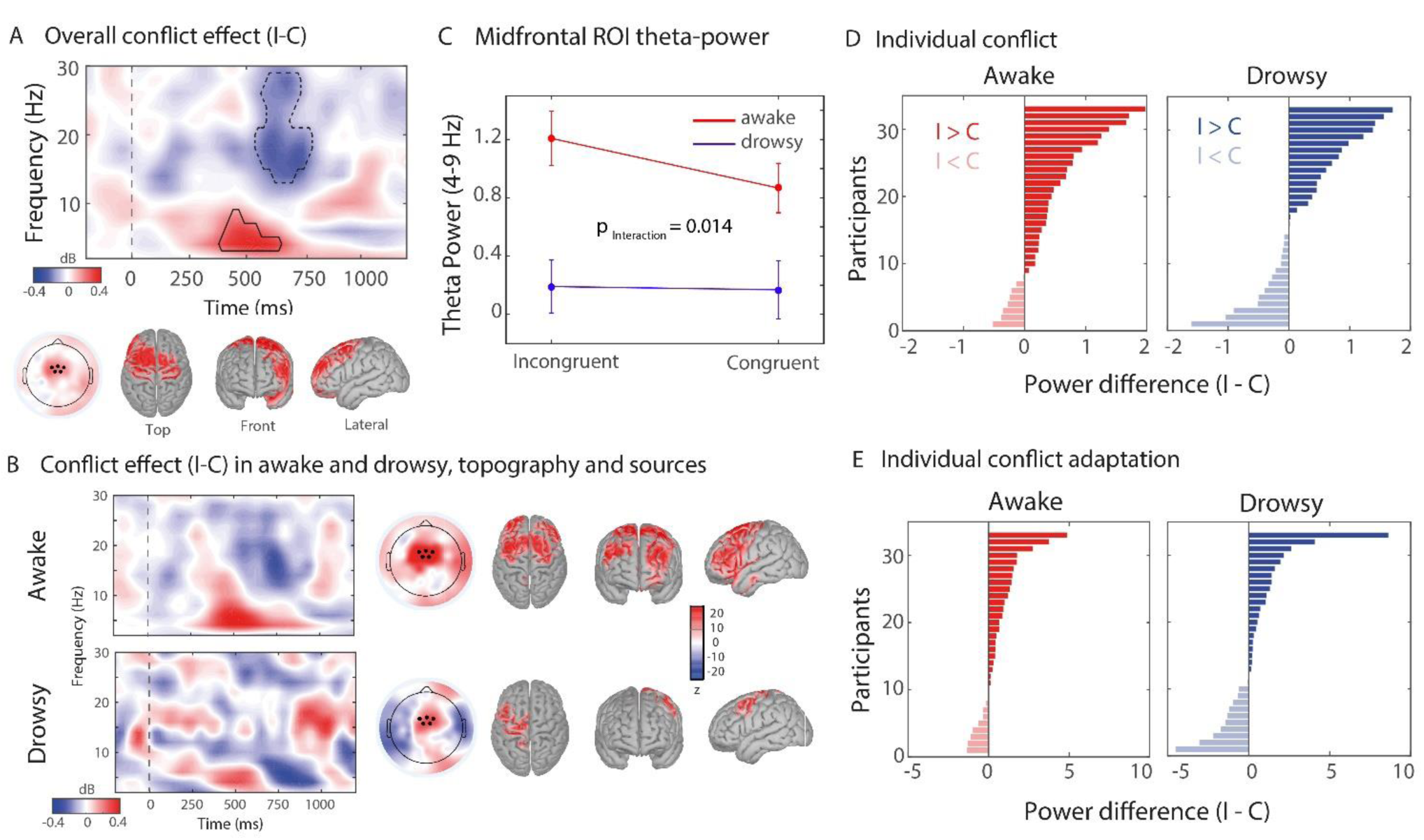
Univariate spectral analysis and sources of midfrontal theta-band oscillations in the awake and drowsy conditions. Conflict effects in terms of time-frequency dynamics across alertness conditions **(A)**, and for the awake versus drowsy states **(B)**, calculated over medial-frontal electrodes **(C)**. **(A)** The black delineated box is the theta-band time-frequency ROI where overall conflict (I-C) was significant over conditions (cluster-based corrected, see Methods). Insets show topographical distribution and sources of oscillatory power within this T-F ROI. Black dots represent the midfrontal EEG electrodes selected for obtaining the conflict-related theta-band power. A source-reconstruction analysis was performed on this time-frequency ROI (z-score) for the overall conflict **(A)** and for awake and drowsy states **(B)**. Activations are depicted on unsmoothed brains; as reconstructed sources were only observed on the surface of the cortex. Sources are for visualization purposes (no statistical testing performed). Group-level **(C)** and individual conflict and individual conflict **(D)** and conflict adaptation effects **(E)** for the awake and drowsy in dB (average ROI power incongruent – average ROI power congruent).

#### Conflict effect in midfrontal (MF) theta-power

First, we evaluated the conflict effect between awake and drowsy trials (I-C) irrespective of previous trial congruency. A RM ANOVA revealed a reliable main effect of alertness (F_1,32_=36.09; p<0.001, 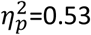, congruency (F_1,32_=6.80; p=0.014, 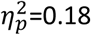, and an interaction between congruency and alertness (F_1,32_=4.63; p=0.014; 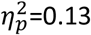, Figure 3C), showing a stronger MF-theta conflict effect in the awake state compared to the drowsy state. Post hoc effects showed higher MF theta for incongruent than congruent trials only in the awake state (awake: t_32_=2.456; p=0.034; drowsy: t_32_=0.305; p=0.761; Tukey corrected for multiple comparisons).

#### Conflict adaptation effect in theta-power

We next evaluated the conflict adaptation effect between awake and drowsy conditions. The extent of conflict related MF theta-power was modulated by previous trial congruency (conflict adaptation: F_1,32_=5.70; p=0.023; 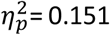, however, no interaction between conflict adaptation and alertness was found (F_1,32_=0.68; p=0.415; 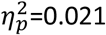 BF_01_=3.21). Finally, as an exploratory analysis, we evaluated the conflict adaptation effect unpacking the results for awake and drowsy conditions separately. In the awake condition, we observed significant conflict adaption in MF theta-band power (F_1,32_=8.47; p=0.007; 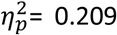 but not in the drowsy state (F_1,32_=2.19; p=0.148; 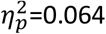 BF_01_=3.84).

In order to visualize the sources of the conflict-related MF theta oscillations observed at the scalp level, we performed source reconstruction analyses, across all conditions (Figure 3A), for the awake and drowsy conflict effects separately (Figure 3B). In line with several fMRI and animal studies performed on awake participants, the conflict-related theta-band signal seems to show hubs in the medial frontal and the dorsolateral prefrontal cortex (Van Veen et al., 2001; Botvinick et al., 2004; Ullsperger et al., 2014), but to a lesser extent in the drowsy condition (Figure 3B).

In addition to the MF theta cluster and in agreement with previous reports (van Gaal et al., 2010; Jiang et al., 2015), an overall conflict effect was observed in the alpha-beta band (cluster p=0.008; frequency range: 13Hz– 29Hz, time range: 580ms–728ms, see encircled region in black, dashed line, in Figure 3A). When trials were split, these results were reliable for the conflict effect in the awake condition (F_1,32_=8.41; p=0.007; 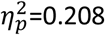 but not for conflict adaptation (F_1,32_=3.24, p=0.081; 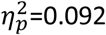, BF_01_=2.021), nor for the drowsy condition in general (conflict effect: F_1,32_=0.05; p=0.488; 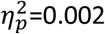, BF_01_=5.252; conflict adaptation: F_1,32_=0.94; p=0.339; 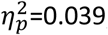, BF_01_=52.135).

#### Relationship between alertness and executive function

We reasoned that testing for differences in congruency before and after stimulus presentation would be helpful to disentangle whether the observed effects are (mainly) related to fluctuations in alertness or potential fluctuations in executive functioning. As a way of disentangling alertness from executive functioning, we directly contrasted the theta-power differences between the pre-trial period used for the alertness classification (−1500 ms to 0 ms pre-stimulus) and the post-stimulus window where conflict was observed (250 ms to 625 ms). A RM ANOVA with the factors time-window and congruency (congruent, incongruent) was performed on the theta-band frequency (frequency range: 4Hz–9Hz, cluster reported in Figure 1A). A significant interaction between window and congruency was observed (F_1,32_=16.919; p<0.001, 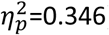, indicating that the post-stimulus theta-band conflict effect was stronger than the pre-stimulus effect. Further, while post hoc analyses confirmed the already-reported difference in congruency in the post-stimulus window (t_32_=2.83, p=0.026), no difference between incongruent and congruent trials was observed in the pre-trial window (t_32_=-1.45, p=0.154; Tukey correct for multiple comparisons). Not enough evidence for a conflict effect in the pre-trial window supports the idea that fluctuations in theta power may index two (partially) separate processes: pre-stimulus theta may mainly reflect fluctuations in alertness levels, whereas post-stimulus theta may signal executive functioning (specifically conflict processing here).

### Multivariate spectral decoding

The hypothesis-driven analysis for the neural signatures focused on the MF theta-band revealed clear conflict detection and conflict adaptation processes for the wake state, but not reliably for the drowsy state. The change of wakefulness in the transition to sleep comes with large changes in neural reconfiguration that might explain this loss of specificity of the neural markers. In order to determine whether a more spatially or temporally extended pattern of neural activity might be underlying the behavioral conflict effect in the drowsy condition observed in behaviour, we performed a spectral Multivariate Pattern Analysis (MVPA) (or ‘spectral decoding’), accounting for possible changes in space, time and frequency of the conflict related neural signatures. Spectral decoding allows one to obtain a measure for the difference between stimuli without having to a priori specify at which electrode(s) or frequency-band this difference emerges, while at the same time picking up subtle differences that might not have been noticed had such an a priori electrode selection been made (Fahrenfort et al., 2018). To do so, we trained classifiers to: (1) distinguish between congruent vs. incongruent trials; (2) distinguish spatial processing in trials where the auditory stimulus was presented from the left vs. the right earbud (i.e. stimulus location); and (3) differentiate trials where the presented auditory stimulus was “left” vs. “right” (i.e. stimulus content). Above-chance classification accuracies imply that relevant information about the decoded stimulus feature is present in the neural data, meaning that some processing of that feature occurred (Hebart and Baker, 2018).

Consistent with the univariate approach for analysing congruency, multivariate decoding showed that information about stimulus congruency was reliably represented in neural data in the awake (Figure 4A), but not in the drowsy state (Figure 4B, p<0.05, cluster-corrected; frequency-range: 2-9 Hz, peak frequency: 6Hz, time-range: 376-810 ms). Assessment of the qualitative difference in the theta-band decoding (4-9 Hz) performance between awake and drowsy states [i.e. an interaction effect: awake AUC values (I-C) minus drowsy AUC values (I-C)). The cluster-based permutation test showed a reliable temporal cluster of increased classifier accuracy for the awake vs. the drowsy condition (p<0.05, cluster-corrected) in the 680-810 ms time-range (Figure 4A right panel). These results are consistent with the interaction effect observed in the univariate analysis performed on the ROI midfrontal theta (Figure 3C).

**Figure 4.**
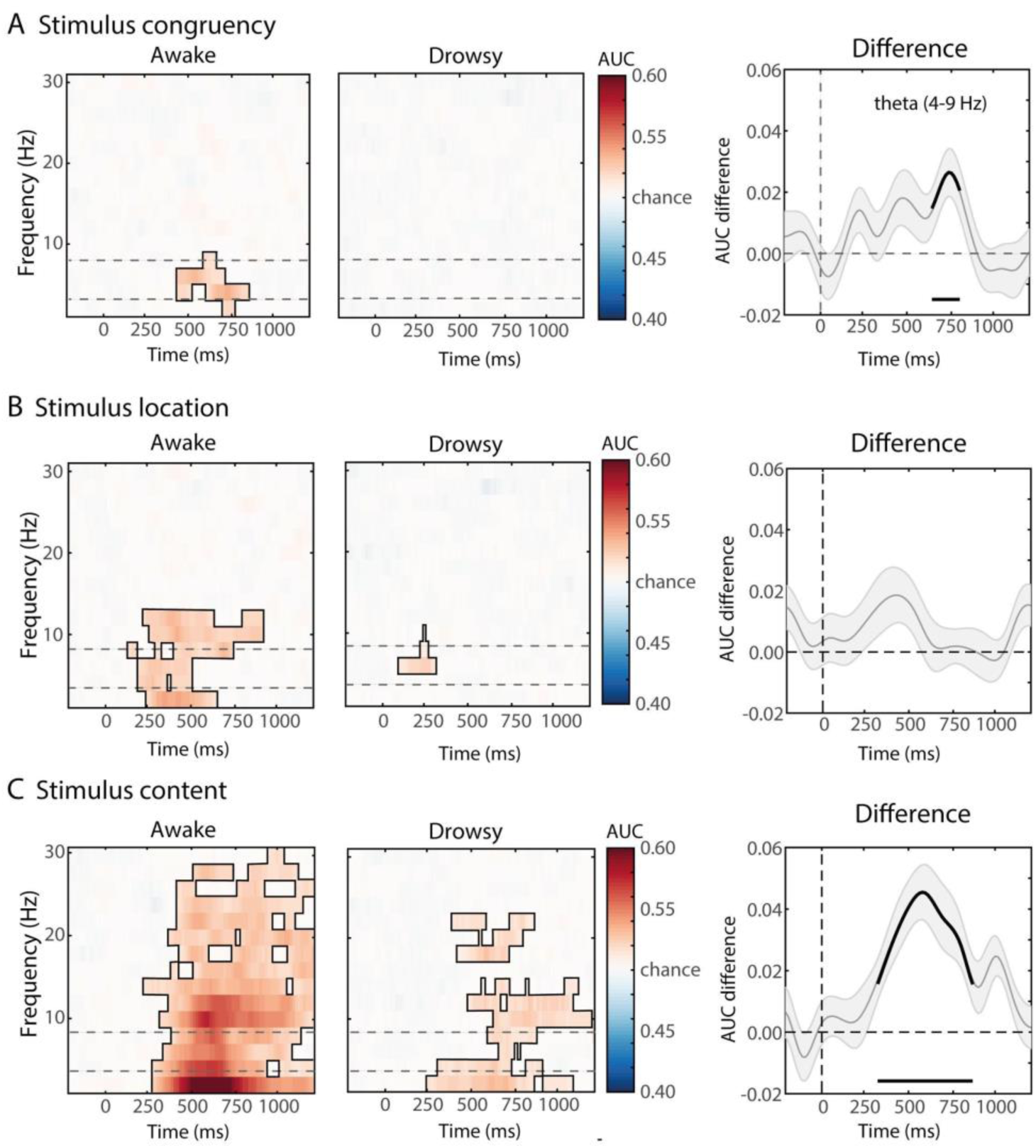
Multivariate spectral decoding of stimuli components in the awake and drowsy condition. Classifier accuracies are depicted across time-frequency charts (2-30 Hz) for the awake and drowsy condition separately, and for the difference between awake and drowsy conditions in the theta-band. Classifier accuracy was thresholded (cluster-based correction, p<0.05) and significant clusters are outlined with a solid black line. In the difference plots on the right, significant differences from chance are highlighted by a black solid line at the bottom of the figures. The dotted lines in the left and middle panel reflect the frequency band used for statistical testing between awake and drowsy states (rightest panels). **(A)** Classifier accuracies for stimulus congruency (“conflict”). Information about congruency was only present in the awake condition. **(B)** Classifier accuracies for stimulus location (“location”). Location of the auditory stimulus could be decoded in both conditions, meaning that information about this stimulus feature is present in both awake and drowsy neural frequency signals. **(C)** Classifier accuracies for stimulus sound identities (“content”). Sound identities of the auditory stimulus could be decoded in both alertness conditions. Differences between awake and drowsy were observed for stimulus congruency and identity but not for stimulus location.

Although the previous analysis revealed that conflict could only be decoded from neural data in the awake state, interestingly, the sound identity and location of the auditory stimuli could be decoded from neural data for both the awake (identity: p<0.001, cluster-corrected, time-range: 240-1200 ms; location: p<0.05, cluster-corrected, time-range: 120-920 ms) and drowsy states (identity: p<0.05, cluster-corrected, time-range: 250-1200 ms; location: p<0.05, cluster-corrected, time-range: 88-300 ms, Figure 4A and Figure 4C). Both content and location showed above chance decoding patterns in the theta-band as well as other frequency bands - depending on the contrast in both awake and drowsy states (Figure 4). This highlights the capacity of the brain to process the semantic and spatial components of the task in parallel under internal modulatory stress (lower arousal). However, the above chance decoding of the individual features forming conflict (location and semantic) but the lack of decoding of conflict itself (congruency), suggest that decreased alertness impacts the capacity of the MFC conflict monitoring system to integrate the semantic and location information required to form conflict.

Summarizing, the above chance performance of the classifiers for low-level stimulus features suggests that location and sound identity were still processed, even during a decreased level of alertness, however, no reliable decoding was found for conflict effects.

### Distributed theta-band information sharing

The fact that a multivariate method analysing the pattern across time, space and frequency did not capture a neural signature of conflict observed behaviourally, suggest a more drastic reconfiguration of the neural processes underlying conflict detection during drowsiness. We reasoned that the neural signatures of conflict may involve changes in connectivity in a wide network of brain regions instead of relatively local power changes. Thus, we hypothesized that a neural metric specifically indexing distributed neural information integration (weighted Symbolic Mutual Information (wSMI); King et al., 2013; Sitt et al., 2014; Imperatori et al., 2019; Canales-Johnson et al., 2020) could in principle capture the conflict effect during drowsiness. wSMI has been shown to capture network reconfiguration both in healthy (Imperatori et al., 2019) and pathological (King et al., 2013; Sitt et al., 2014) states of alertness. Importantly, due its sensitivity to highly non-linear coupling (Imperatori et al., 2019), wSMI has been useful for tracking the neural information dynamics underlying perceptual content formation (Canales-Johnson et al., 2020). Thus, we performed this analysis as a possible post-hoc hypothesis for the reconfiguration of the underlying networks supporting cognitive control. The wSMI can be calculated at different time-scales, which can be implemented by the variable tau (see Methods). Here we used a tau of 32 ms (~4-9 Hz) and therefore the wSMI measure captures non-linear information integration in the theta-band domain. The time window for theta-band wSMI was calculated on the significant time window observed in the spectral contrast of Figure 3A (380-660 ms). A RM ANOVA revealed a reliable main effect of alertness (F_1,32_=56.10; p<0.001, 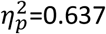, which is in line with previous results showing increased wSMI in states associated with high levels, vs. low levels, of arousal and consciousness (King et al., 2013; Sitt et al., 2014; Imperatori et al., 2019). More interestingly, we also observed an interaction between congruency and alertness for long-distance wSMI in the theta-band (F_1,32_=5.50; p=0.025; 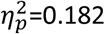, Figure 5A). Post hoc tests showed higher wSMI for incongruent than congruent trials only in the drowsy state (drowsy: t_32_=2.456; p=0.034; awake: t_32_=0.305; p=0.761; Tukey corrected for multiple comparisons). Individual differences in theta-band wSMI for each participant in the awake (right) and drowsy (left) conditions are shown in Figure 5B.

**Figure 5.**
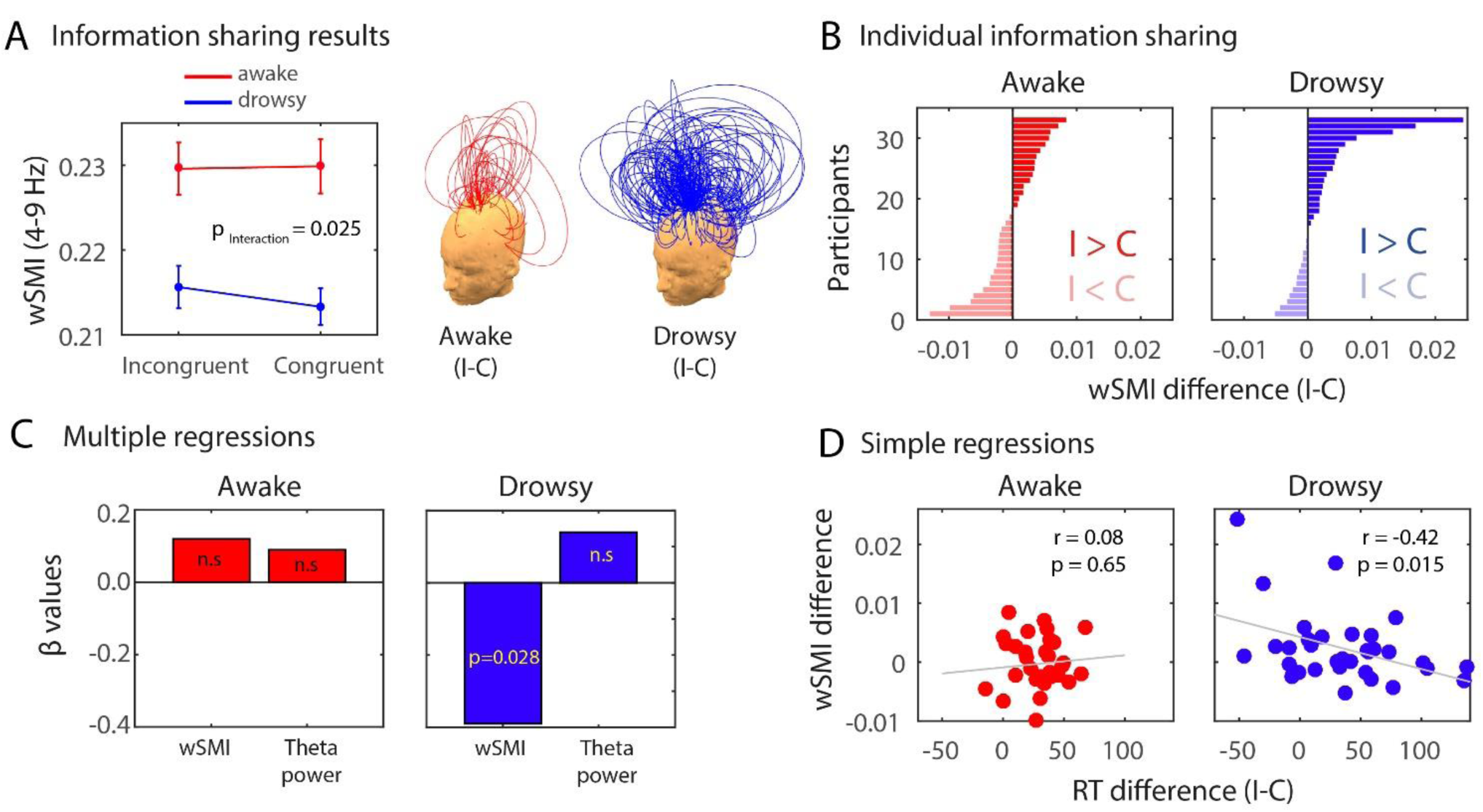
Long-distance theta-band information sharing during conflict in awake and drowsy. **(A)** Long-distance wSMI in the theta-band during the conflict effect. Each arc represents a functional connection between a pair of electrodes, and the height of the arc represents the value of the wSMI difference for that pair (incongruent-congruent; awake condition in green and drowsy condition in magenta). Theta-band wSMI was calculated between each midfrontal ROI electrode (shown in Figure 3) and every other electrode outside the ROI. wSMI values within the midfrontal ROI were discarded from the analyses since we aimed at evaluated information integration between distant electrode pairs. **(B)** Individual differences in theta-band wSMI for each participant in the awake (right) and drowsy (left) conditions. **(C)** Beta coefficients for two separate multiple regressions using RT difference (I-C) as predicted variable and theta power difference (I-C) and wSMI difference (I-C) as regressors **(D)** Pearson’s correlation for awake and drowsy conditions between RT differences and wSMI difference.

In order to further control for potential wSMI effects in other frequency ranges traditionally associated with cognitive control, we computed alpha-band wSMI (tau of 24 ms; ~10-14 Hz, 380 ms - 660 ms) and beta-band wSMI (tau of 16 ms; ~15-20 Hz, same time window as beta-power effect of Figure 3A: 580 ms – 728 ms). In the case of the alpha-band wSMI, a main effect of alertness (F_1,32_=89.86; p<0.001; 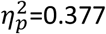, but no main effect of congruency (F_1,32_=0.02; p=0.874) nor an interaction between alertness and congruency (F_1,32_=0.618; p=0.437) was observed. Similarly, a main effect of alertness (F_1,32_=52.51; p<0.001; 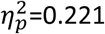 but no main effect of congruency (F_1,32_=1.08; p=0.307) nor interaction (F_1,32_=2.48; p=0.125) were observed for the beta-band wSMI. These results further suggest the specificity of theta-band wSMI in the neural reconfiguration of cognitive control during diminished alertness.

### Brain-behaviour relationships

We further investigated, in an exploratory manner, the statistical dependencies between information integration in the theta-band, information sharing (wSMI) and the strength of the behavioural conflict effect. Separate multiple regressions were performed on the awake and drowsy states, using as regressors the MF theta-power ROI differences (I-C) and the distributed theta-wSMI differences (I-C) (Figure 5C). The conflict effect (RT difference: I-C) was used as the predicted variable. In the drowsy condition model (R^2^=0.20; F2,30=3.68; p=0.037), distributed theta-wSMI predicted the conflict effect in RT (β=-0.39; p=0.028), while MF theta was not a reliable predictor (β=0.14; p=0.665). On the other hand, in the awake condition model (R^2^=0.02; F2,30=; p=0.613), none of the regressors predicted the conflict effect significantly (MF theta: β=0.12; p=0.514; distributed theta-wSMI: β=0.09; p=0.609). This relationship was also described in terms of a simple Pearson’s correlation, showing a significant correlation between RT difference and wSMI difference for the drowsy (r=-0.42; p=0.015) but not for the awake condition (r=0.08; p=0.65) (Fig 5D). These results show that the distributed information, but possibly not the local spectral power in the same neural signal (theta-band), underlies the behavioural conflict effect observed in the drowsy state.

## DISCUSSION

In this article we describe the impact of changes in our arousal state during conflict detection and conflict adaptation processes during normal waking fluctuations. We found the expected behavioural manifestations of decreased alertness, namely higher group variability in RTs and slower RTs when people were drowsy as compared to actively awake (Lal and Craig, 2001; Huang et al., 2009; Goupil and Bekinschtein, 2012; Bareham et al., 2014; Comsa et al., 2019). Further, we observed reliable conflict effects with decreased alertness. However, contrary to recent observations (Gevers et al. (2015), we observed conflict adaptation even in the drowsy state, despite participants’ decreased alertness (Figure 2). The effects of conflict (current trial) and conflict adaptation (trial-by-trial) were reliable independently of the states of alertness (see Figure 2 for individual participant’s data), suggesting a spared capacity to resolve conflict arising from the incongruity between the meaning and the side of the world where the word was presented. These arousal modulations on the capacity of executive control have been found in fatigued participant and after sleep deprivation (Gevers et al., 2015), but others showed only slower reaction times and no Stroop Effects (Sagaspe et al., 2006; Cain et al., 2011; Bratzke et al., 2012). However, we show here that normal fluctuations of arousal in well-rested participants yield no strong detrimental effects in the resolution of conflict. In short, humans still experience conflict while mildly drowsy, and even if they react slower, they respond to incongruity in a similar manner as when fully awake and attentive.

Although conflict processing was relatively maintained in behavioural terms, its neural signatures changed. The principles of neural reorganization are a much debated topic in neuroscience (Dahmen and King, 2007; Shine et al., 2019) but there is agreement in the flexibility of the brain networks to maintain or preserve psychological function in the face of insult, and/or internal or external modulatory factors (Siuda-Krzywicka et al., 2016; Singh et al., 2018). Here we found a dissociation between the behavioural and neural markers of conflict and conflict adaptation effect with the change in alertness. The classic conflict-induced theta-band power changes were no longer reliable during low alertness while slower RT for incongruent trials remained. Furthermore, multivariate whole-brain analyses showed convergent results with the univariate approach. These findings suggest that the changes exerted by the diminished alertness elicited a reconfiguration of the brain networks putatively responsible for the neural resolution of the conflict, resulting in the disappearance of the theta power difference in the conflict contrast and rise of a more distributed network in the same frequency band, as evidenced by the increased information shared in theta connectivity.

The networks implicated in cognitive control are not only supported by correlations with brain activity, but also by causal interventions. In rodents, a dissociation has been proposed between prefrontal cortices in the causal support of control functions in which the dorso-medial prefrontal cortex seems to be implicated in memory for motor responses. This includes response selection and the temporal processing of information, whereas ventral regions of the medial prefrontal cortex seem implicated in interrelated ‘supervisory’ attentional functions, including attention to stimulus features and task contingencies (or action–outcome rules), attentional set-shifting, and behavioural flexibility (Dalley et al., 2004). In humans, causal evidence is sparse due to a scarcity of patients with specific (frontal) lesions. However, the impairment of simple cognitive control and trial-by-trial influence is shown in a small but well-structured study in which dorsal anterior cingulate cortex (dACC) damage disrupted the ability to make an adaptive choice between actions (but not stimuli) following a win on the previous trial. Moreover, orbitofrontal (OFC) damage similarly disrupted choices between stimuli, but not actions (Camille et al., 2011). Furthermore, in a large (n=344) correlational study Gläscher et al. (2012) found that impairments in cognitive control (response inhibition, conflict monitoring, and switching) was associated with dorsolateral prefrontal cortex and anterior cingulate cortex lesions. These medial prefrontal areas that have been proposed as the origin of the theta power modulation in conflict tasks are thus causally implicated in cognitive control and lend further support for the search of other correlates that would capture the conflict effect during an arousal challenge. Our results show the involvement of the medial prefrontal cortex and, to a lesser degree, the lateral prefrontal cortex in the classic conflict Simon task in agreement with the literature. However, we observed a more distributed network supporting the resolution of conflict, when internal resources lose efficiency as a consequence of a drowsy state (Comsa et al., 2019).

Another framework to interpret our results is provided by Posner’s taxonomy of attention (Posner, 2008). The framework proposes three brain networks that contribute to attention: alerting, orienting, and executive control. In our project we assess the interplay between the alerting network, achieving and maintaining a state of high sensitivity to stimuli, and executive control, the mechanisms for resolving conflict. Under this framework the fluctuations in alertness induced by the different task settings in our experiment (awake vs drowsy session) pertain to the maintenance of optimal vigilance and we study how decreases in alertness impact the system dedicated to conflict resolution. Other studies have looked at the interaction between the alerting system and executive attentional systems and found either no interacting effects or limited interaction between systems (Fan et al., 2002; Fossella et al., 2002) in healthy participants. However, in patients, the modulation of the alerting network improved cognitive control (Robertson et al., 1995; Sturm et al., 2006). We interpret our findings as an indication that fluctuations in the alerting network impact the cognitive control network. and the neural reconfiguration that we have observed is a reflection of neural resilience and of compensatory mechanisms that the human brain instantiates when high levels of performance have to be maintained when challenged by arousal changes.

The cognitive processes leading up to conflict experience involve the extraction of meaning (“left” or “right”) and location from where the stimulus came from. Thus, if the two factors are congruent (“left” coming from the left side of space) conflict is supposed to be absent and the participant responses fast, but when the word comes from the other side of the space (“left” presented in the right side of space) conflict arises and the responses slow down, reflecting further processes necessary to resolve conflict. We hypothesised that the specific perceptual and semantic component of location and content, respectively, would be decodable in the spectral domain as the participants responded correctly to the stimuli. Both content and location showed above chance decoding patterns in the theta-band as well as other frequency bands-depending on the contrast in both awake and drowsy states (Figure 4). This highlights the capacity of the brain to process the semantic and spatial components of the task in parallel under internal modulatory stress (lower arousal). In order to capture the integration between these two components by cognitive control networks, we looked for decodability of conflict in the spectral domain (stimulus congruency). The patterns showed the expected theta-band power difference (restricted to theta) only in the alert state, the lack of decodable patterns in the frequency space further suggested that the neural signatures of the conflict resolution would be found somewhere else, pointing to connectivity as a possible candidate.

Since we knew that there is strong evidence that neural markers of conflict can be found in brain signals, we decided to turn to information sharing under three premises. First, a neural measure of information sharing could in principle capture directly the information integration between stimulus content and stimulus location necessary for generating the conflict effect in our task. Second, the dynamic nature of neural information integration (Imperatori et al., 2019; Canales-Johnson et al., 2020) may be able to capture the reconfiguration of neural networks during the transition from an alert to a drowsy state of mind. Finally, as the reorganization of networks could be reflected in the need for larger information capacity of the brain when challenged (by drowsiness), the measure chosen can be conceptually framed as deriving from a computational principle. Although cortical reorganization with age and after insult have been extensively studied, the cognitive flexibility, or “cognitive fragmentation” resulted from an internally generated change –drowsiness has hardly been captured (Goupil and Bekinschtein, 2012).

We would also like to note that by manipulating alertness levels (by having an awake and a drowsy session) we cannot fully rule out that other cognitive functions may potentially fluctuate alongside. For example, it may be that arousal fluctuations partially go hand in hand with fluctuations in sustained attention (Foucher et al., 2004) and/or levels of proactive control (i.e. the sustained and anticipatory maintenance of goal-relevant information) (Braver, 2012). We have aimed to isolate alertness fluctuations as accurately as possible by using an algorithm that separates awake from drowsy trials that has previously been validated in different domains (Jagannathan et al., 2018) and by focusing on physiological measures (e.g. alpha power, the presence of ripples, vertex sharp waves, etc.) that are firmly rooted in a long history of work on alertness (arousal) fluctuations (Hori, 1985; Goupil and Bekinschtein, 2012). Further, we have provided support for the idea that changes in theta power can reflect changes in multiple processes, because control analyses revealed that theta power indexes separate processes before (i.e. alertness levels) and after (i.e. cognitive control) stimulus presentation. In future work it would be valuable though to further isolate unique influences of fluctuations in alertness, (sustained) attention and proactive control on (reactive) cognitive control processes, such as conflict detection/resolution (Braver, 2012).

These methods of tackling the system during transitions could be conceptually regarded as causal if the processes at play (wakefulness and cognitive control) are viewed as partially independent. The case of drowsiness as a causal mode in cognitive neuroscience may prove to be very useful in the exploration of how cognition is fragmented or remains resilient under (reversible) perturbations of wakefulness (Bareham et al., 2014; Kouider et al., 2014; Comsa et al., 2019). However, it cannot be regarded as causal in the sense of perturbing the system externally as TMS, TDCs or pharmacology would do, since wakefulness is naturally occurring and intimately interrelated with cognitive processes.

One possible explanation for the call for wider networks to resolve conflict during drowsiness would be the need for involvement of extended neural resources to solve the same task, as seen previously in older adults when they are matched in performance to younger adults (Reuter-Lorenz and Cappell, 2008; Spreng et al., 2017). Convergent evidence is drawn from cognitive control studies, where the frontoparietal control networks are further recruited with higher cognitive load (Liang et al., 2016; Fransson et al., 2018), tasks possibly reflecting the higher need for neural resources. In other words, the brain’s capacity for plasticity allows for the expansion of conflict networks in cases where another element in the system (e.g. drowsiness) draws resources away (internal challenge) from the neural systems typically underlying cognitive control.

In conclusion, we have shown that although participants were generally sluggish, the conflict effect reflected behaviourally in slower responses to conflicting information compared to non-conflicting information was still intact during a drowsy state of alertness. The changes in the neural signatures of conflict from local theta oscillations (awake condition) to a long-distance distributed theta network (drowsy condition) suggests a relative reconfiguration of the underlying neural processes subserving cognitive control when the system is affected by lower levels of alertness.

## Acknowledgements

We would like to thank Dr. Sridhar Rajan Jagannathan for his technical assistance and Lavazza for its unconditional support. The research leading to these results was supported by the Wellcome Trust Biomedical Research Fellowship WT093811MA awarded to TAB, core funds from the Department of Psychology at Cambridge

## References

Allefeld C, Görgen K, Haynes JD (2016) Valid population inference for information-based imaging: From the second-level t-test to prevalence inference. Neuroimage 141:378–392.

Baillet S, Mosher JC, Leahy RM (2001) Electromagnetic brain mapping. IEEE Signal Process Mag 18:14–30.

Bareham CA, Manly T, Pustovaya O V, Scott SK, Bekinschtein TA (2014) Losing the left side of the world: rightward shift in human spatial attention with sleep onset. Sci Rep 4:5092 Available at: http://www.pubmedcentral.nih.gov/articlerender.fcgi?artid=4035582&tool=pmcentrez&rendertype=abstract.

Bekinschtein T, Cologan V, Dahmen B, Golombek D (2009) You are only coming through in waves: wakefulness variability and assessment in patients with impaired consciousness. Prog Brain Res.

Borbély AA, Daan S, Wirz-Justice A, Deboer T (2016) The two-process model of sleep regulation: A reappraisal. J Sleep Res.

Botvinick MM, Cohen JD, Carter CS (2004) Conflict monitoring and anterior cingulate cortex: An update. Trends Cogn Sci 8:539–546.

Bratzke D, Steinborn MB, Rolke B, Ulrich R (2012) Effects of sleep loss and circadian rhythm on executive inhibitory control in the stroop and simon tasks. Chronobiol Int 29:55–61.

Braver TS (2012) The variable nature of cognitive control: A dual mechanisms framework. Trends Cogn Sci.

Cai W, Chen T, Ryali S, Kochalka J, Li CSR, Menon V (2016) Causal Interactions Within a Frontal-Cingulate-Parietal Network During Cognitive Control: Convergent Evidence from a Multisite-Multitask Investigation. Cereb Cortex 26:2140–2153.

Cain SW, Silva EJ, Chang AM, Ronda JM, Duffy JF (2011) One night of sleep deprivation affects reaction time, but not interference or facilitation in a Stroop task. Brain Cogn 76:37–42.

Camille N, Tsuchida A, Fellows LK (2011) Double dissociation of stimulus-value and action-value learning in humans with orbitofrontal or anterior cingulate cortex damage. J Neurosci 31:15048–15052.

Canales-Johnson A, Billig AJ, Olivares F, Gonzalez A, Garcia M del C, Silva W, Vaucheret E, Ciraolo C, Mikulan E, Ibanez A, Huepe D, Chennu S, Bekinschtein TA (2020) Dissociable Neural Information Dynamics of Perceptual Integration and Differentiation during Bistable Perception. Cereb Cortex.

Cavanagh JF, Frank MJ, Klein TJ, Allen JJB (2010) Frontal theta links prediction errors to behavioral adaptation in reinforcement learning. Neuroimage 49:3198–3209.

Cohen MX, Donner TH (2013) Midfrontal conflict-related theta-band power reflects neural oscillations that predict behavior. J Neurophysiol 110:2752–2763.

Cohen MX, Ridderinkhof KR (2013) EEG Source Reconstruction Reveals Frontal-Parietal Dynamics of Spatial Conflict Processing. PLoS One 8.

Cohen MX, Ridderinkhof KR, Haupt S, Elger CE, Fell J (2008) Medial frontal cortex and response conflict: Evidence from human intracranial EEG and medial frontal cortex lesion. Brain Res 1238:127–142.

Cohen MX, van Gaal S (2014) Subthreshold muscle twitches dissociate oscillatory neural signatures of conflicts from errors. Neuroimage 86:503–513.

Comsa IM, Bekinschtein TA, Chennu S (2019) Transient Topographical Dynamics of the Electroencephalogram Predict Brain Connectivity and Behavioural Responsiveness During Drowsiness. Brain Topogr 32:315–331.

Dahmen JC, King AJ (2007) Learning to hear: plasticity of auditory cortical processing. Curr Opin Neurobiol 17:456–464.

Dalley JW, Cardinal RN, Robbins TW (2004) Prefrontal executive and cognitive functions in rodents: Neural and neurochemical substrates. In: Neuroscience and Biobehavioral Reviews, pp 771–784.

Desimone R, Duncan J (1995) Neural Mechaisms of Selective Attention. Annu Rev Neurosci 18:193–222.

Driel J van, Olivers CNL, Fahrenfort JJ (2019) High-pass filtering artifacts in multivariate classification of neural time series data. bioRxiv:530220 Available at: https://www.biorxiv.org/content/10.1101/530220v2.

Egner T, Hirsch J (2005) Egner, T., & Hirsch, J. (2005). Cognitive control mechanisms resolve conflict through cortical amplification of task-relevant information. Nature neuroscience, 8(12), 1784–90. doi:10.1038/nn1594Cognitive control mechanisms resolve conflict through cortica. Nat Neurosci 8:1784–1790 Available at: http://www.ncbi.nlm.nih.gov/pubmed/16286928.

Fahrenfort JJ, van Driel J, van Gaal S, Olivers CNL (2018) From ERPs to MVPA using the Amsterdam Decoding and Modeling toolbox (ADAM). Front Neurosci 12.

Fan J, McCandliss BD, Sommer T, Raz A, Posner MI (2002) Testing the efficiency and independence of attentional networks. J Cogn Neurosci 14:340–347.

Fossella J, Sommer T, Fan J, Wu Y, Swanson JM, Pfaff DW, Posner MI (2002) Assessing the molecular genetics of attention networks. BMC Neurosci 3.

Foucher JR, Otzenberger H, Gounot D (2004) Where arousal meets attention: A simultaneous fMRI and EEG recording study. Neuroimage.

Fransson P, Schiffler BC, Thompson WH (2018) Brain network segregation and integration during an epoch-related working memory fMRI experiment. Neuroimage 178:147–161.

Gevers W, Deliens G, Hoffmann S, Notebaert W, Peigneux P (2015) Sleep deprivation selectively disrupts top-down adaptation to cognitive conflict in the Stroop test. J Sleep Res 24:666–672.

Gläscher J, Adolphs R, Damasio H, Bechara A, Rudrauf D, Calamia M, Paul LK, Tranel D (2012) Lesion mapping of cognitive control and value-based decision making in the prefrontal cortex. Proc Natl Acad Sci U S A 109:14681–14686.

Goupil L, Bekinschtein TA (2012) Cognitive processing during the transition to sleep. Arch Ital Biol 150:140–154.

Gramfort A, Papadopoulo T, Olivi E, Clerc M (2010) OpenMEEG: Opensource software for quasistatic bioelectromagnetics. Biomed Eng Online 9.

Gratton G, Coles MGH, Donchin E (1992) Optimizing the Use of Information: Strategic Control of Activation of Responses. J Exp Psychol Gen 121:480–506.

Gunzelmann G, Gluck KA, Price S, Van Dongen HPA, Dinges DF (2012) Decreased Arousal as a Result of Sleep Deprivation: The Unraveling of Cognitive Control. In: Integrated Models of Cognitive Systems.

Harrison Y, Horne JA, Rothwell A (2000) Prefrontal Neuropsychological Effects of Sleep Deprivation in Young Adults—a Model for Healthy Aging? Sleep 23:1–7.

Hebart MN, Baker CI (2018) Deconstructing multivariate decoding for the study of brain function. Neuroimage 180:4–18.

Hori T (1985) Spatiotemporal changes of eeg activity during waking-sleeping transition period. Int J Neurosci.

Huang RS, Jung TP, Makeig S (2009) Tonic changes in EEG power spectra during simulated driving. In: Lecture Notes in Computer Science (including subseries Lecture Notes in Artificial Intelligence and Lecture Notes in Bioinformatics), pp 394–403.

Imperatori LS, Betta M, Cecchetti L, Canales-Johnson A, Ricciardi E, Siclari F, Pietrini P, Chennu S, Bernardi G (2019) EEG functional connectivity metrics wPLI and wSMI account for distinct types of brain functional interactions. Sci Rep 9.

Jagannathan SR, Ezquerro-Nassar A, Jachs B, Pustovaya O V., Bareham CA, Bekinschtein TA (2018) Tracking wakefulness as it fades: Micro-measures of alertness. Neuroimage 176:138–151.

Jiang J, Correa CM, Geerts J, van Gaal S (2018) The relationship between conflict awareness and behavioral and oscillatory signatures of immediate and delayed cognitive control. Neuroimage 177:11–19.

Jiang J, Zhang Q, van Gaal S (2015) Conflict awareness dissociates theta-band neural dynamics of the medial frontal and lateral frontal cortex during trial-by-trial cognitive control. Neuroimage 116:102–111.

King JR, Sitt JD, Faugeras F, Rohaut B, El Karoui I, Cohen L, Naccache L, Dehaene S (2013) Information sharing in the brain indexes consciousness in noncommunicative patients. Curr Biol 23:1914–1919.

Kouider S, Andrillon T, Barbosa LS, Goupil L, Bekinschtein TA (2014) Inducing task-relevant responses to speech in the sleeping brain. Curr Biol 24:2208–2214.

Lal SKL, Craig A (2001) A critical review of the psychophysiology of driver fatigue. Biol Psychol 55:173–194.

Liang X, Zou Q, He Y, Yang Y (2016) Topologically Reorganized Connectivity Architecture of Default-Mode, Executive-Control, and Salience Networks across Working Memory Task Loads. Cereb Cortex 26:1501–1511.

Luu P, Tucker DM, Makeig S (2004) Frontal midline theta and the error-related negativity: Neurophysiological mechanisms of action regulation. Clin Neurophysiol 115:1821–1835.

Maris E, Oostenveld R (2007) Nonparametric statistical testing of EEG- and MEG-data. J Neurosci Methods 164:177–190.

Miller EK, Cohen JD (2001) An Integrative Theory of Prefrontal Cortex Function. Annu Rev Neurosci 24:167–202.

Nigbur R, Cohen MX, Ridderinkhof KR, Stürmer B (2012) Theta dynamics reveal domain-specific control over stimulus and response conflict. J Cogn Neurosci 24:1264–1274.

Pastötter B, Dreisbach G, Bäuml KHT (2013) Dynamic adjustments of cognitive control: Oscillatory correlates of the conflict adaptation effect. J Cogn Neurosci 25:2167–2178.

Posner MI (2008) Measuring alertness. In: Annals of the New York Academy of Sciences, pp 193–199.

Ratcliff R, Van Dongen HPA (2011) Diffusion model for one-choice reaction-time tasks and the cognitive effects of sleep deprivation. Proc Natl Acad Sci U S A 108:11285–11290.

Reuter-Lorenz PA, Cappell KA (2008) Neurocognitive aging and the compensation hypothesis. Curr Dir Psychol Sci 17:177–182.

Robbins TW (1996) Dissociating executive functions of the prefrontal cortex. Philos Trans R Soc B Biol Sci 351:1463–1471.

Robertson IH, Tegnér R, Tham K, Lo A, Nimmo-Smith I (1995) Sustained Attention Training for Unilateral Neglect: Theoretical and Rehabilitation Implications. J Clin Exp Neuropsychol 17:416–430.

Sagaspe P, Sanchez-Ortuno M, Charles A, Taillard J, Valtat C, Bioulac B, Philip P (2006) Effects of sleep deprivation on Color-Word, Emotional, and Specific Stroop interference and on self-reported anxiety. Brain Cogn 60:76–87.

Shine JM, Breakspear M, Bell PT, Ehgoetz Martens K, Shine R, Koyejo O, Sporns O, Poldrack RA (2019) Human cognition involves the dynamic integration of neural activity and neuromodulatory systems. Nat Neurosci 22:289–296.

Singh AK, Phillips F, Merabet LB, Sinha P (2018) Why Does the Cortex Reorganize after Sensory Loss? Trends Cogn Sci 22:569–582.

Sitt JD, King JR, El Karoui I, Rohaut B, Faugeras F, Gramfort A, Cohen L, Sigman M, Dehaene S, Naccache L (2014) Large scale screening of neural signatures of consciousness in patients in a vegetative or minimally conscious state. Brain 137:2258–2270.

Siuda-Krzywicka K, Bola Ł, Paplińska M, Sumera E, Jednoróg K, Marchewka A, Śliwińska MW, Amedi A, Szwed M (2016) Massive cortical reorganization in sighted braille readers. Elife 5.

Spreng RN, Shoemaker L, Turner GR (2017) Executive Functions and Neurocognitive Aging. In: Executive Functions in Health and Disease, pp 169–196.

Stroop JR (1935) Studies of interference in serial verbal reactions. J Exp Psychol 18:643–662.

Sturm W, Thimm M, Küst J, Karbe H, Fink GR (2006) Alertness-training in neglect: Behavioral and imaging results. Restor Neurol Neurosci 24:371–384.

Swick D, Ashley V, Turken U (2011) Are the neural correlates of stopping and not going identical? Quantitative meta-analysis of two response inhibition tasks. Neuroimage 56:1655–1665.

Tadel F, Baillet S, Mosher JC, Pantazis D, Leahy RM (2011) Brainstorm: A user-friendly application for MEG/EEG analysis. Comput Intell Neurosci 2011.

Trujillo LT, Allen JJB (2007) Theta EEG dynamics of the error-related negativity. Clin Neurophysiol 118:645–668.

Tucker AM, Whitney P, Belenky G, Hinson JM, Van Dongen HPA (2010) Effects of sleep deprivation on dissociated components of executive functioning. Sleep 33:47–57.

Ullsperger M, Danielmeier C, Jocham G (2014) Neurophysiology of performance monitoring and adaptive behavior. Physiol Rev 94:35–79.

van Gaal S, Lamme VAF, Ridderinkhof KR (2010) Unconsciously triggered conflict adaptation. PLo S One 5.

Van Veen V, Cohen JD, Botvinick MM, Stenger VA, Carter CS (2001) Anterior cingulate cortex, conflict monitoring, and levels of processing. Neuroimage 14:1302–1308.

